# Non-muscle Myosin II acts as a negative feedback mediator to control cell contraction dynamics

**DOI:** 10.1101/2025.09.12.675762

**Authors:** Carolin Gierse, Jennifer Hanemann, Leif Dehmelt

## Abstract

Local cell contraction dynamics play a crucial role in tissue and cell morphogenesis. Contractions near the cell edge drive highly dynamic cell shape changes during cell migration and contraction pulses in central cell attachment areas are involved in mechanotransduction. Previously, we identified a signal network in adherent mammalian cells, that generates mechanosensitive contraction pulses, in which the cell contraction regulator Rho is controlled by fast positive feedback amplification via GEF-H1, and by a slow negative feedback that depends on actomyosin activity. However, the precise mechanism of this negative feedback in adherent cells, in particular if it is mediated via actin or Myosin-based components, was still unclear. Here, using numerical simulations of this system, we predicted that the cell contraction signal network dynamics are strongly inhibited both by inhibition and by constitutive activation of the actomyosin component Myosin-II. We confirmed these predictions experimentally by direct inhibition of Myosin-II and by activation via constitutively active ROCK1. Furthermore, constitutive activation of Myosin-II leads to an accumulation of Myosin-II next to the nuclei which spatially correlated with a corresponding shift of Rho activity dynamics from the cell center to the cell edge, showing that constant Myosin activation can spatially restrict cell contraction dynamics. Finally, light-induced rapid recruitment of ROCK1 to the plasma membrane strongly activated and recruited Myosin-II, and at the same time depleted Actin and inhibited Rho activity at the plasma membrane. We conclude that negative feedback in the cell contraction signal network of adherent mammalian cells is mediated by Myosin-II, and that actin does not act as the predominant inhibitor in this system.

## Introduction

It is well established that cells process both chemical and mechanical signals to modulate their function. For example, stem cells can be induced to differentiate into distinct cell types purely by differences in the elasticity of their surrounding matrix (ECM) (Engler et al., 2006). To sense such differences in substrate elasticity, cells have to convert mechanical signals into biochemical signals in a process that is called mechanotransduction (Aguilar-Cuenca et al., 2014). Cells utilize a complex, multilevel system to perform this task. Important components of this system are integrin transmembrane receptors that link extracellular structures to the cytosol, and adapter proteins in specialized adhesion complexes that connect these receptors with contractile actomyosin structures. In several model organisms, highly dynamic actomyosin-driven cell contraction pulses were observed (Martin et al., 2009; Bement et al., 2015; Graessl et al., 2017; Nalbant and Dehmelt, 2018; Saha et al., 2018; Kim et al., 2018; Michaux et al., 2018), which were proposed to act in an active sensing process to transduce extracellular mechanical signals more efficiently compared to constant tension (Cui et al., 2015; Graessl et al., 2017; Nalbant and Dehmelt, 2018). Indeed, Rho activity pulses were observed in differentiating stem cells, and pulse frequency was modulated by substrate elasticity (Sampayo et al., 2023). Furthermore, such pulses can also play an instructive role, as it was shown that optogenetically-controlled oscillatory Rho activity was sufficient to steer stem cell differentiation (Sampayo et al., 2023). However, the molecular details of the underlying mechanism of this active sense of touch for cells are still unclear.

Previous work in adherent mammalian cells revealed key features of the signal network that can generate mechanosensitive pulses of cell contraction (Graessl et al., 2017). In particular, this previous work revealed an activator-inhibitor signal network topology, in which the small GTPase RhoA rapidly amplifies its activity by recruiting its activator GEF-H1 to the plasma membrane, and in which this amplification is inhibited by a slower actomyosin-dependent process (Graessl et al., 2017). Such a combination of fast positive and slow negative feedback is known to generate pulsatile, oscillatory or excitable system dynamics (Novák and Tyson, 2008), and the general signal network topology of this system was confirmed by combining theoretical and experimental approaches(Kamps et al., 2020). In particular, these studies clearly demonstrated the molecular mechanism for positive feedback, however, the molecular details on negative feedback were still poorly understood.

Strikingly, while work in adherent mammalian cells supported a specific role for contractile actomyosin structures and non-muscle Myosin 2 in negative feedback, investigations in other systems, including *Xenopus* and Starfish oocytes or *C. elegans* embryos suggested a more general involvement of filamentous actin and found that non-muscle Myosin 2 was dispensable for the generation of pulsatile signal network dynamics (Nishikawa et al., 2017; Michaux et al., 2018). More detailed investigations revealed that in *Xenopus* and Starfish oocytes, F-actin recruits the RhoGAP RGA3/4 to the cell cortex, where it can inhibit Rho activity to close a negative feedback loop that controls cell contraction pulses (Michaud et al., 2022), and a similar mechanism drives negative feedback regulation in cell contraction pulses of *C. elegans* embryos (Michaux et al., 2018).

Interestingly, such an F-actin based Rho recruitment mechanism can also be enforced in adherent mammalian cells by overexpression of the actin-associated RhoGAP Myo9b, however, the frequency of the generated pulses was much higher compared to pulses that are generated at normal Myo9b levels, and their amplitude was much lower (Graessl et al., 2017). In contrast, the high amplitude, low frequency pulses were proposed to be generated by a non-muscle Myosin II mediated feedback mechanism (Graessl et al., 2017).

Molecularly, the inhibitory function of active Myosin could be mediated by its interaction with the catalytic DH domain of the feedback mediator GEF-H1 (Lee et al., 2010; Kamps et al., 2020). This interaction could compete with GEF-H1 mediated Rho activation and thereby disrupt positive feedback amplification of Rho (Lee et al., 2010; Kamps et al., 2020). This idea is supported by the observation that Myosin recruitment to the plasma membrane is anti-phasic to Rho activity. Furthermore, direct inhibition of Myosin or inhibition of the Myosin activating Rho effector kinase ROCK results in a strong reduction of pulsatile network activity in adherent cells. However, these manipulations will also indirectly affect F-actin structures. Therefore, the nature of the inhibitory mechanism in adherent cells was still unclear, in particular with regard to the overwhelming evidence that an F-actin-based negative feedback mechanism controls cell contraction dynamics in *Xenopus* and starfish oocytes and in *C. elegans* embryos.

Here, we directly investigated the role of Myosin II in the control of cell contraction dynamics in mammalian adherent cells. In particular, we applied both long-term and rapid perturbations in the cell contraction signal network by manipulating the function of the Myosin activating kinase ROCK. Long-term perturbations were based on the overexpression of constitutively active (CA-)ROCK1 and resulted in a strong inhibition of cell contraction network dynamics. Rapid perturbations via photochemically-controlled plasma membrane targeting of ROCK1 resulted in a rapid increase in Myosin, rapid decrease in actin and strong reduction of Rho activity dynamics at the plasma membrane. From these investigations we conclude that the delayed negative feedback in the mechanosensitive cell contraction pulses of adherent mammalian cells is specifically mediated by Myosin II.

## Results and Discussion

To investigate the molecular mechanism that drives pulsatile cell contraction dynamics, we capitalized on our previous work, in which we established that either ectopic expression of microtubule binding deficient GEF-H1 C53R (Krendel et al., 2002) or nocodazole treatment to release GEF-H1 from microtubules (Krendel et al., 2002; Chang et al., 2008) can strongly stimulate cell contraction signal network dynamics. Furthermore, GEF-H1 plasma membrane recruitment dynamics are closely correlated with Rho activity in space and time due to the direct interaction between GEF-H1 and active Rho (Kamps et al., 2020). Therefore, GEF-H1 was used in this study either to stimulate the dynamics of the cell contraction signal network, to detect local Rho activity patterns, or both.

Figure 1A-C and Movie_Figure1 show a representative cell that generates the low frequency, high amplitude cell contraction pulses of active Rho/GEF-H1 and Myosin IIa, which we typically observe in this system. Similar to previous observations (Graessl et al., 2017), we observe anti-phasic plasma membrane recruitment kinetics and a time delay of about 30 seconds that separates the active Rho/GEF-H1 and Myosin IIa pulses (Figure 1C) (Graessl et al., 2017; Kamps et al., 2020). As shown previously, these pulses of Rho and Myosin IIa plasma membrane recruitment were associated with local centripetal flow of actomyosin structures, showing that Myosin transiently and locally activated within each pulse (Graessl et al., 2017).

**Figure 1:**
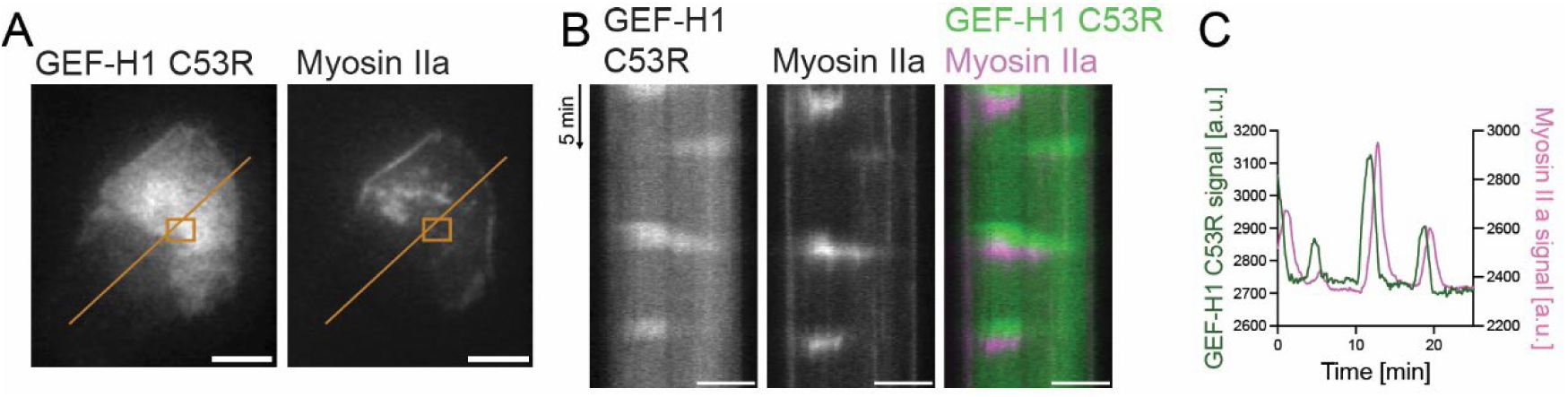
Pulsatile activity dynamics in the cell contraction signal network of adherent mammalian cells. A-C: Analysis of GEF-H1/Myosin II plasma membrane recruitment dynamics, which are linked to their activity state under control of the cell contraction signal network. A: Representative total internal reflection fluorescence microscopy (TIRF-M) images of mCitrine-GEF-H1 C53R and mTurquoise-Myosin IIa. B: Kymographs corresponding to orange lines in B. C: Quantification of GEF-H1 C53R and Myosin IIa signals corresponding to orange boxes in A. Scale bars: 10 µm.

### Constitutive, non-feedback inhibition or activation of Myosin decreases pulsatory cell contraction dynamics

To gain a more quantitative understanding of the negative feedback in this system, we used a stochastic differential equation (SDE) system that describes the dynamics of the key system components GEF-H1, active Rho, and Myosin. This system was developed previously based on experimental data (Kamps et al., 2020) and a simplified scheme of the underlying biochemical reactions (Figure 2A). We first aimed to validate this SDE system by comparing simulations with experiments that we already performed previously (Figure 2B and Figure S1) and implemented additional parameters into this SDE system (see Methods for details) to consider the following experimental manipulations: 1) A reduction of the total concentration of Myosin to represent Myosin inhibition via the small molecule inhibitor blebbistatin. 2) A reduction of the effective rate constant that links Rho activity to Myosin activation to represent the inhibition of the Rho effector ROCK via the small molecule inhibitor Y276732.

**Figure 2:**
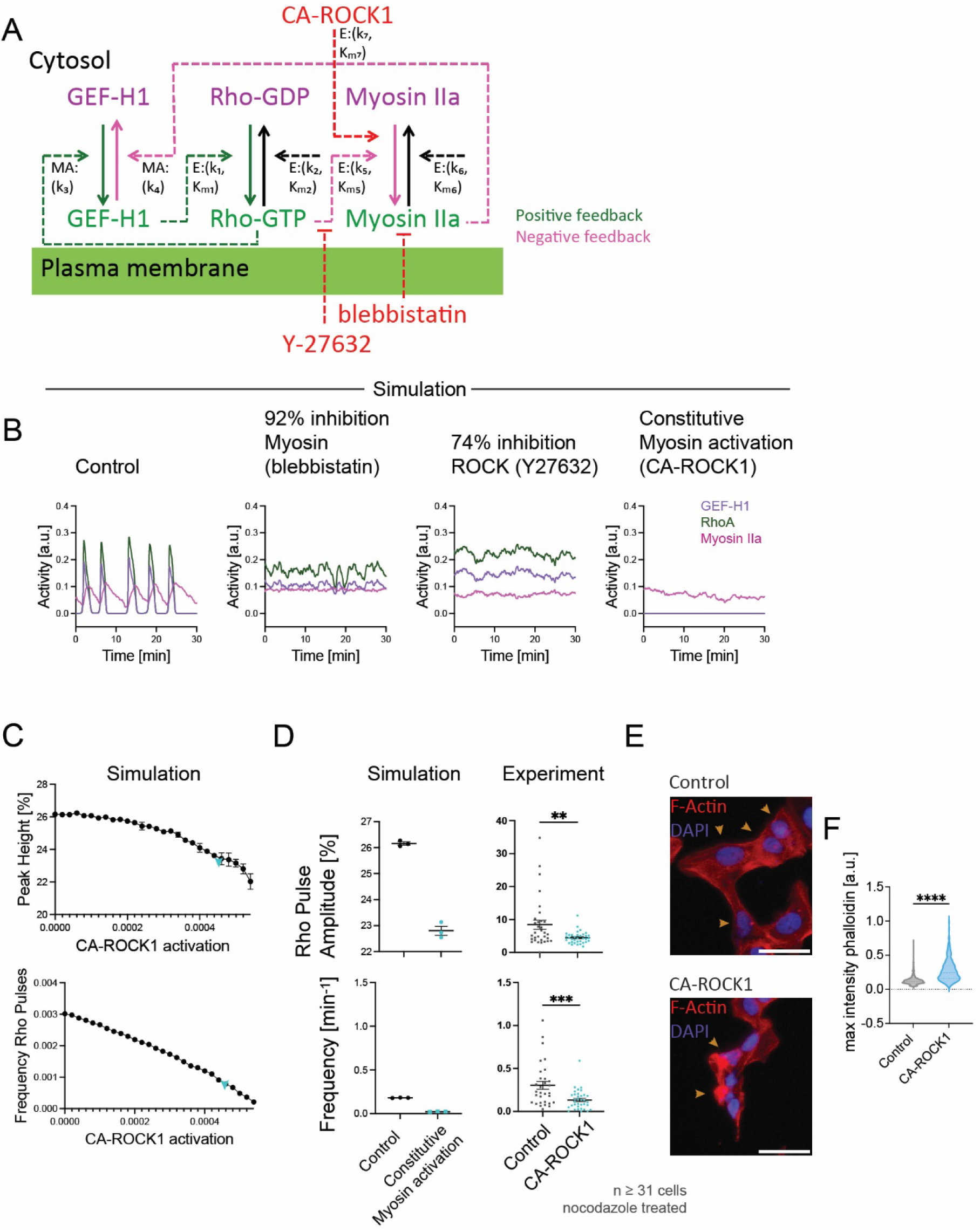
Constitutive inhibition or activation of Myosin IIa inhibits the cell contraction signal network dynamics of adherent mammalian cells. A: Simplified scheme for biochemical reactions that were proposed to generate oscillatory and excitable dynamics of cell contraction (see Methods for details; MA denotes reactions via mass action, E denotes enzymatic reactions; dotted lines represent causal links that are mediated by biochemical reactions, solid lines denote changes in the activty state that are associated with translocation between the cytosol and plasma membrane). B: SDE simulations of GEF-H1 stimulated signal network dynamics in the absence or presence of the Myosin inhibitor blebbistatin (corresponding to 92% inhibition) (see also Figure S1A), the ROCK inhibitor Y27632 (corresonding to 74% inhibition) (see also Figure S1B) or constitutively active ROCK1 (CA-ROCK1 activation level corresponding to k_7_=0.0006, see also panels C-D). All experiments and simulations shown in this study are based on a stimulated signal network state, which was experimentally achieved either by expression of constitutively active GEF-H1 or nocodazole-mediated release of microtubule-sequested endogenous GEF-H1, and included in simulations by increasing the total concentration of available, active GEF-H1 (G_T_=0.3, see Methods for details). C: Quantification of Rho activity peak amplitude or pulse frequency as a function of increasing CA-ROCK1 concentrations obtained via SDE simulations (n=3 independent simulations for each value). CA-ROCK1 activation in the x-axis corresponds to the rate constant k_7_ in the scheme shown in panel A. D: Comparison of Rho activity peak amplitude or pulse frequency obtained via SDE simulations (left) or experimental analysis (right) in the absence or presence of CA-ROCK1. Experimental analysis was performed by measuring Rho activity dynamics in U2OS cells using a Rhotekin-based translocation sensor in U2OS cells, which co-expressed either an mCherry-CA-ROCK1 fusion protein, or the mCherry fluorophore alone as control. Signal network dynamics were stimulated by the addition of nocodazole (30 µM). SDE simulations were performed at a concentration of CA-ROCK1 that corresponds to the blue arrowheads in C. E: Representative wide-field images of mCherry-CA-ROCK1 or mCherry control expressing cells, which were stained for F-actin (atto488-phalloidin) and DNA/nuclei (Hoechst 33342). Orange arrowheads indicate transfected cells that express the respective construct. F: Quantification of the maximal F-actin signal corresponding to the conditions shown in E. (n=3 independent repetitions with ≥ 694 cells). Error bars represent standard error of the mean. **: p < 0.01, ***: p < 0.001, ****: p < 0.0001, Student’s t-Test, Scale bars: 50 µm.

Interestingly, moderate Myosin II inhibition had very little effect on pulse dynamics in our simulations, however, if Myosin was inhibited by more than ≈92%, dynamics slowed down dramatically and quickly approached very long periods. These results agree with experimental results from Graessl et al. (2017), in which only very high concentrations of blebbistatin (Myosin inhibitor, 30 µM) inhibited or reduced pulsatile Rho activity dynamics in cells and this is likely due to the very high endogenous expression levels of non-muscle Myosin II in our cell system (Beck et al., 2011). Inhibition of ROCK by Y27632 also inhibited signal network dynamics in our SDE simulations, however, compared to Myosin inhibition via blebbistatin, we observed a more gradual, concentration-dependent reduction of peak frequency and peak height. This effect in also in agreement with our previously published experimental data on peak frequency (Graessl et al., 2017), and peak amplitude (Melanie Graessl). Thus, our simulations agree with previous experimental data and therefore support the general biochemical rationale behind the SDE system and specifically the idea that negative feedback is mediated by Myosin II.

Next, we introduced constitutively active ROCK1 (CA-ROCK1) as a new component into the SDE system, to predict how a continuous, Rho activity-independent activation of Myosin II would affect signal network dynamics (Figure 2A, see Methods for details). Simulations based on this extended SDE system predicted that even low levels of constitutive activation of Myosin via CA-ROCK1 would strongly inhibit network dynamics in a dose-dependent manner (Figure 2B-D). This prediction based on our theory was confirmed experimentally in U2OS cells: Expression of CA-ROCK1 indeed strongly suppressed Rho activity dynamics (Figure 2D). This effect of CA-ROCK1 could be explained by the uncontrolled, Rho-independent stimulation of Myosin, which would then inhibit GEF-H1 (Lee et al., 2010) and prevent its ability to amplify Rho activity (Graessl et al., 2017).

To confirm that our CA-ROCK1 construct is functional, we measured its effect on F-actin structures in U2OS cells. Based on previous studies (Ishizaki et al., 1996; Leung et al., 1996), we expected that the constitutive and continuous activation of Myosin would lead to the abundant formation of contractile actomyosin structures and a local accumulation of F-actin structures. Indeed, we find that CA-ROCK1 leads to the formation of prominent, strongly stained F-actin foci (Figure 2E), and quantification of the maximal F-actin signals (Figure 2F) and F-actin signal standard deviation (A) using cell profiler confirmed this observation. Taken together, these observations show that Rho-independent, constitutive, non-feedback activation or inhibition of Myosin inhibits Rho activity dynamics in adherent, mammalian cells. This suggests that Myosin is not merely a functional output of an autonomous system that only acts upstream, but instead an integral component of the cell contraction signal network and is compatible with the idea that it mediated negative feedback regulation.

### Constitutive Myosin activation leads to local Myosin accumulation and local inhibition of cell contraction signal network activity

We noticed that the strongly stained F-actin foci stimulated by CA-ROCK (Figure 2E, bottom) were typically localized to more central cell regions near the nucleus. This was in contrast to control cells, in which F-actin was relatively evenly distributed in the whole cell attachment area (Figure 2E, top). F-actin foci are typically generated by contractile actomyosin forces and thus are expected to contain high levels of Myosin II. We therefore wondered if these local foci have a local or a global effect on Rho activity dynamics.

To investigate this question, we co-transfected CA-ROCK1 (Figure 3A, right) or the empty parental plasmid (Figure 3A, left) together with fluorescently labelled Myosin IIa heavy chain to characterize Myosin IIa dynamics under these conditions in living cells. In the control condition, the majority of cells (>90%) show typical Myosin II structures near the plasma membrane, in particular stress fibers as well as small patches that were associated with dynamic cell contraction pulses (Figure 3A-C). However, less than 20% of CA-ROCK1 expressing cells contained normal Myosin structures (Figure 3A-C), and the majority (>70%) of the cells formed dense accumulations of Myosin II (Figure 3A-C), which were considerably less dynamic compared to cell contraction pulses in control conditions. In some cells, a flow of Myosin II towards the cell center can be observed (see for example the kymograph in Figure 3A, right).

**Figure 3:**
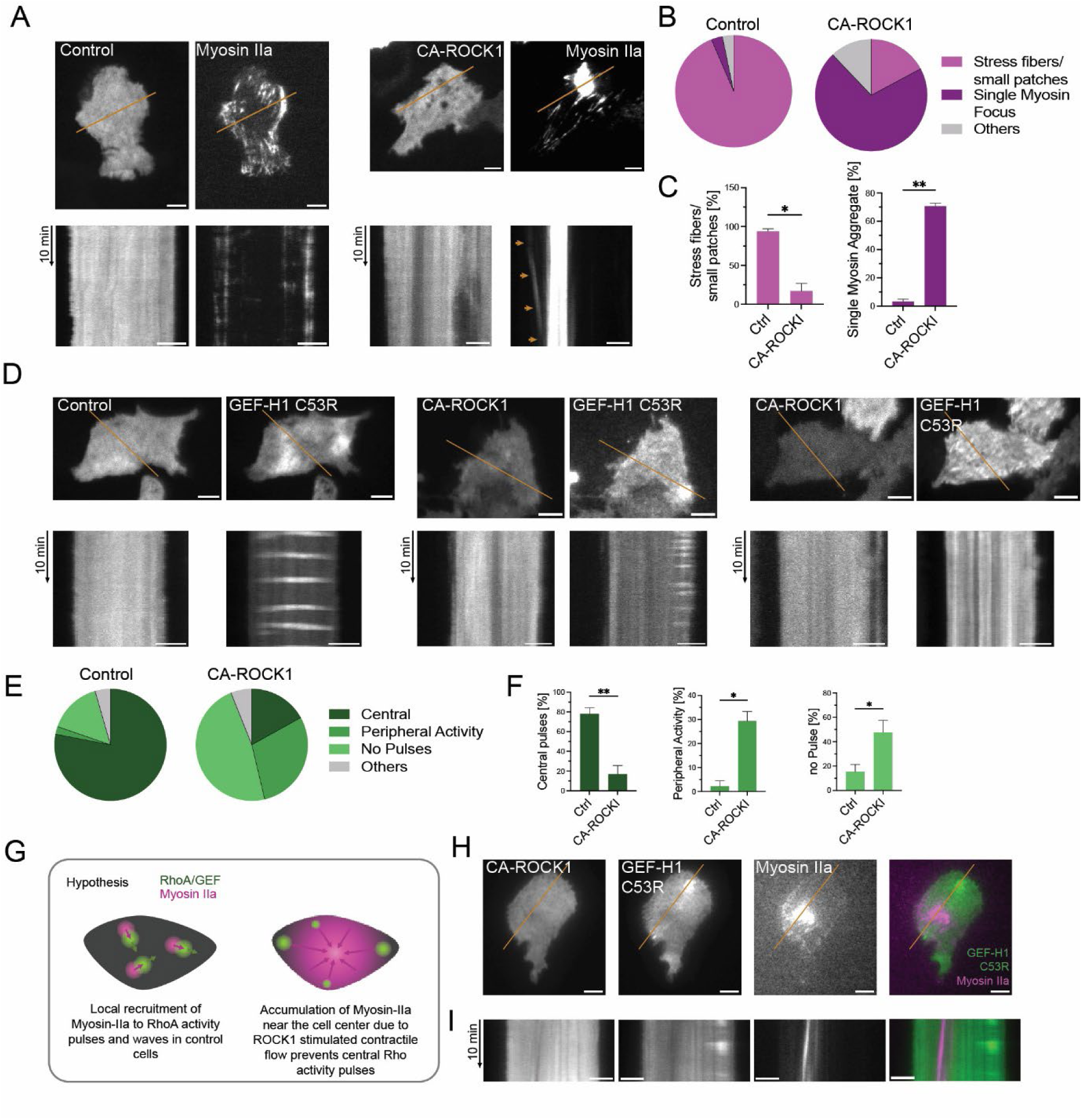
Continuous activation of Myosin IIa via constitutively active ROCK1 restricts cell contraction signal network dynamics to peripheral cell attachment areas. A-C: Analysis of Myosin IIa response to long-term CA-ROCK1 expression. A: Top: Representative TIRF images of cells expressing mCherry as control (left) or mCherry-CA-ROCK1 (right) together with mCitrine-non-muscle Myosin heavy chain IIa (Myosin IIa). Bottom: Kymographs corresponding to orange lines in TIRF images (top). Orange arrows indicate retrograde flow of Myosin IIa towards the single Myosin IIa focus near the cell center. B-C: Quantitative analysis of the effect of CA-ROCK1 on Myosin IIa structures in cells. D-F: Analysis of the effect of long-term CA-ROCK1 expression on cell contraction signal network dynamics. D: Top: Representative TIRF images of cells co-expressing mCherry as control (left) or mCherry-CA-ROCK1 (middle and right) and mCitrine-GEF-H1 C53R. Bottom: Kymographs corresponding to orange lines in TIRF images (top). E-F: Quantitative analysis of the effect of CA-ROCK1 on the spatio-temporal dynamics of GEF-H1 C53R. G: Proposed mechanism of spatio-temporal shift of GEF-H1 C53R dynamics in cells treated with CA-ROCK1. Constitutive activation of Myosin leads to the formation of a single Myosin focus near the cell center and a contractile flow that depletes active Myosin from the cell periphery. Based on this hypothesis, the inhibitory action of Myosin on Rho activity would be reduced in the cell periphery, and thus would restrict pulsatile signal network dynamics to peripheral cell areas. H-I: Experimental support for the hypothesis proposed in G. H: Representative TIRF images of cells co-expressing mCherry-CA-ROCK1, mCitrine-GEF-H1 C53R, and mTurqoise-non-muscle Myosin heavy chain IIa. I: Kymographs corresponding to regions indicated in H (orange lines). n = 3 independent experiments with >42 cells for A-C and >32 cells for D-F. Error bars represent standard error of the mean. *: p < 0.05, **: p < 0.01, Student’s t-Test. Scale bars: 10 µm.

Next, we investigated how GEF-H1 C53R expressing cells with increased Rho/GEF-H1 activity dynamics respond to CA-ROCK1 co-expression. As shown in Figure 3D-F, we found that about 80% of control cells that lack CA-ROCK1 generate Rho/GEF-H1 activity pulses in central cell attachment areas, similar to the typical observations shown in Figure 1. In contrast, only about 20% of the CA-ROCK1 co-transfected cells show this typical activity pattern (Figure 3D-F). Instead, a substantial fraction of these cells (29%) exhibited Rho/GEF-H1 activity pulses exclusively in a narrow region of the peripheral cell attachment area, while only 2% of cells lacking CA-ROCK1 showed this particular spatio-temporal pattern (Figure 3D-F).

Because CA-ROCK1 activates Myosin continuously and without feedback control, we hypothesized that the resulting constitutive activation of Myosin leads to an increased and continuous centripetal actomyosin flow from the periphery of the cell towards more central regions of the cell attachment area. In the cell center, increased Myosin could specifically inhibit Rho amplification, while peripheral cell regions are spared from the continuous presence of high Myosin concentrations due to its removal via centripetal flow. This hypothetical mechanism could explain the observed shift of pulses from central to more peripheral areas, which are then less affected by increased local Myosin levels (Figure 3G). Indeed, simultaneous imaging of the dynamics of active Rho/GEF-H1 and Myosin confirmed this idea (Figure 3H-I). Rho activity dynamics were absent in the proximity of the Myosin focus and only observed at most distant regions of the cell periphery. Interestingly, Rho activity was clearly reduced not only within regions of the highest Myosin concentration, but also in regions around the Myosin focus, which had weaker, but nevertheless detectable Myosin signals (Figure 3I). Taken together, we conclude that Rho-independent activation of ROCK1 leads to Myosin II accumulation near the nucleus with a constant centripetal flow from the periphery towards the cell center, and that the associated Myosin concentration gradient at the plasma membrane inhibits dynamic Rho activity in central cell areas, only allowing activity in the periphery, where Myosin is depleted by its self-generated contractile flow.

### Rapid activation of Myosin reduces Rho activity pulse dynamics

The experiments presented above support the idea that Myosin activity acts as an inhibitor of Rho activity dynamics. However, long-term, continuous activation downstream of CA-ROCK1 could lead to various changes in cell state. For example, CA-ROCK1 can phosphorylate and thereby directly activate the formin FHOD1, which then could lead to increased F-actin polymerization (Takeya et al., 2008). To investigate the immediate sequence of events downstream of active ROCK1 in more detail, we introduced rapid photochemically-controlled perturbations of ROCK1 at the plasma membrane of living cells. First, we employed the previously established molecular activity painting (MAP, Figure 4A) method (Chen et al., 2017; Kamps et al., 2020) to recruit ROCK1 to a small region at the plasma membrane upon local uncaging of a photo-chemical dimerizer (Figure 4B-D). Interestingly, plasma membrane recruitment of full length, wild-type ROCK1 was sufficient to strongly recruit and activate Myosin II in the corresponding cell region, suggesting that increasing the local concentration of this molecule is sufficient to stimulate its downstream effects (Figure 4B-D).

**Figure 4:**
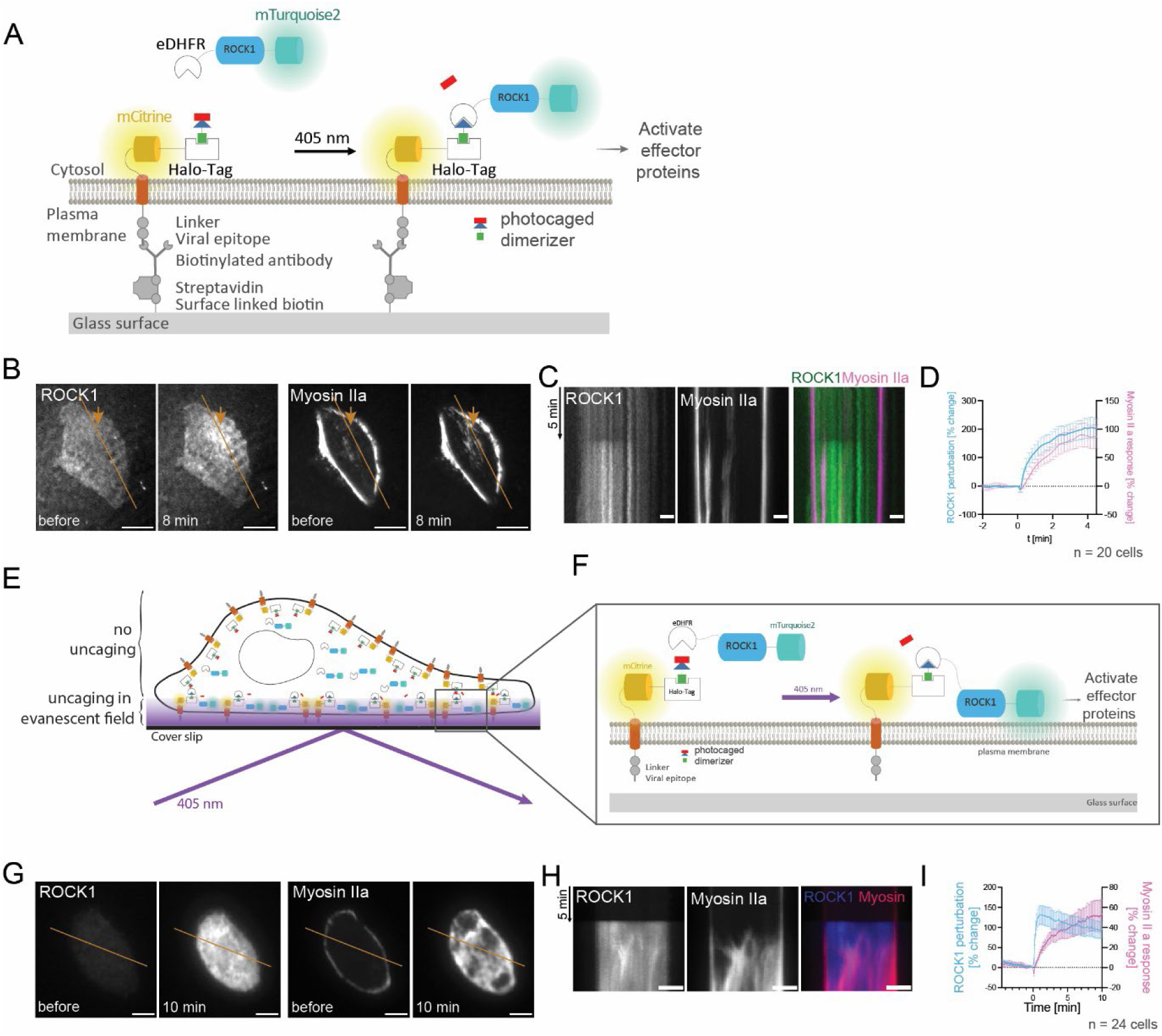
Activation of Myosin by light-controlled plasma membrane recruitment of ROCK1. A: Schematic of the original Molecular Activity Painting approach used to locally stimulate Myosin asctivity in cells. In this method, a single pulse of near UV light (405nm) is used to uncage a photochemical dimierizer. This uncaging enables the interaction of eDHFR and a plasma membrane anchored Halo-Tag, leading to the plasma membrane recruitment and enrichment of ROCK1 fused to eDHFR. This enrichment of ROCK1 can locally activate Myosin at the plasma membrane. Total internal reflection fluorescence microscopy (TIRF-M) is used to measure the kinetics of the ROCK1 perturbation and of the downstream Myosin response. B: Representative TIRF image of a U2OS cell that co-expresses mTurquoise2-ROCK1-eDHFR (to introduce the perturbation, left) and mCherry-non-muscle Myosin heavy chain IIa (Myosin IIa, for response readout, right). Orange arrows indicate the uncaging position. C: Kymographs corresponding to orange lines in B. D: Quantification of the mTurquoise2-ROCK1-eDHFR perturbation and mCherry-non-muscle Myosin heavy chain IIa (Myosin IIa) response (n = 20 cells from 3 independent experiments). E-F: Schematic of a modified Molecular Activity Painting approach used to stimulate Myosin activity in the entire cell attachment area of cells. In the original method (A), uncaging was performed by near UV light (405nm) illumination in a single, diffraction limited spot. In contrast, this variant uses a single pulse of near UV TIRF illumination to selectively uncage photochemical dimerizes only within the evanescent field of the TIRF microscope, i.e. within about 100nm distance from the cell attachement surface. This enables the plasma membrane recruitment and enrichment of ROCK1 to the plasma membrane not only at a local spot, but instead in the whole cell attachment area. Total internal reflection fluorescence microscopy is again used to measure the kinetics of the ROCK1 perturbation and of the downstream Myosin IIa response. In contrast to the original method, the strong immobilization of the Halo-Tag to prevent lateral diffusion is not necessary to achieve strong perturbation that are stable for several minutes. G: Representative TIRF images of chemo-optogenetic mTurquoise2-ROCK1-eDHFR plasma membrane recruitment and response of mCherry-non-muscle Myosin heavy chain IIa (Myosin IIa). H: Kymographs corresponding to orange lines in G. I: Quantification of co-recruitment of Myosin IIa construct (n = 24 cells from 3 independent experiments). Error bars represent standard error of the mean. Scale bars: 10 µm.

To investigate the effect of signaling downstream of ROCK1 on Rho activity dynamics, we further optimized this rapid perturbation by: 1) Uncaging the dimerizer in the entire cell attachment area via UV light using TIRF illumination (Figure 4E) and 2) by avoiding additional manipulations for immobilization of the artificial receptor, which is critical for highly local perturbations, but not required for the envisioned perturbation in a larger area of the plasma membrane (Figure 4F). As shown inFigure 4G-I, this variant allowed a very effective, and very fast, switch-like perturbation of ROCK1 within the entire cell attachment area. This switch-like ROCK1 perturbation was accompanied by a very strong Myosin IIa response that started immediately and increased continuously over several minutes.

Next, we performed this rapid perturbation in the presence of the Rhotekin-based Rho activity sensor after stimulation of Rho activity dynamics with nocodazole (Figure 5A-G). Strikingly, ROCK1 recruitment resulted in an immediate drop of the basal Rho sensor signals by about 5% (Figure 5D). Furthermore, a quantitative analysis of the Rho activity dynamics revealed a substantial decrease in both the pulse frequency and amplitude, as well as the standard deviation of the Rho sensor signal (Figure 5G). In contrast, in samples that were not treated with the dimerizer we did not observe any change in the Rho signals, demonstrating that the observed effects are specific to the plasma membrane recruitment of ROCK1 (Figure 5E-G).

**Figure 5:**
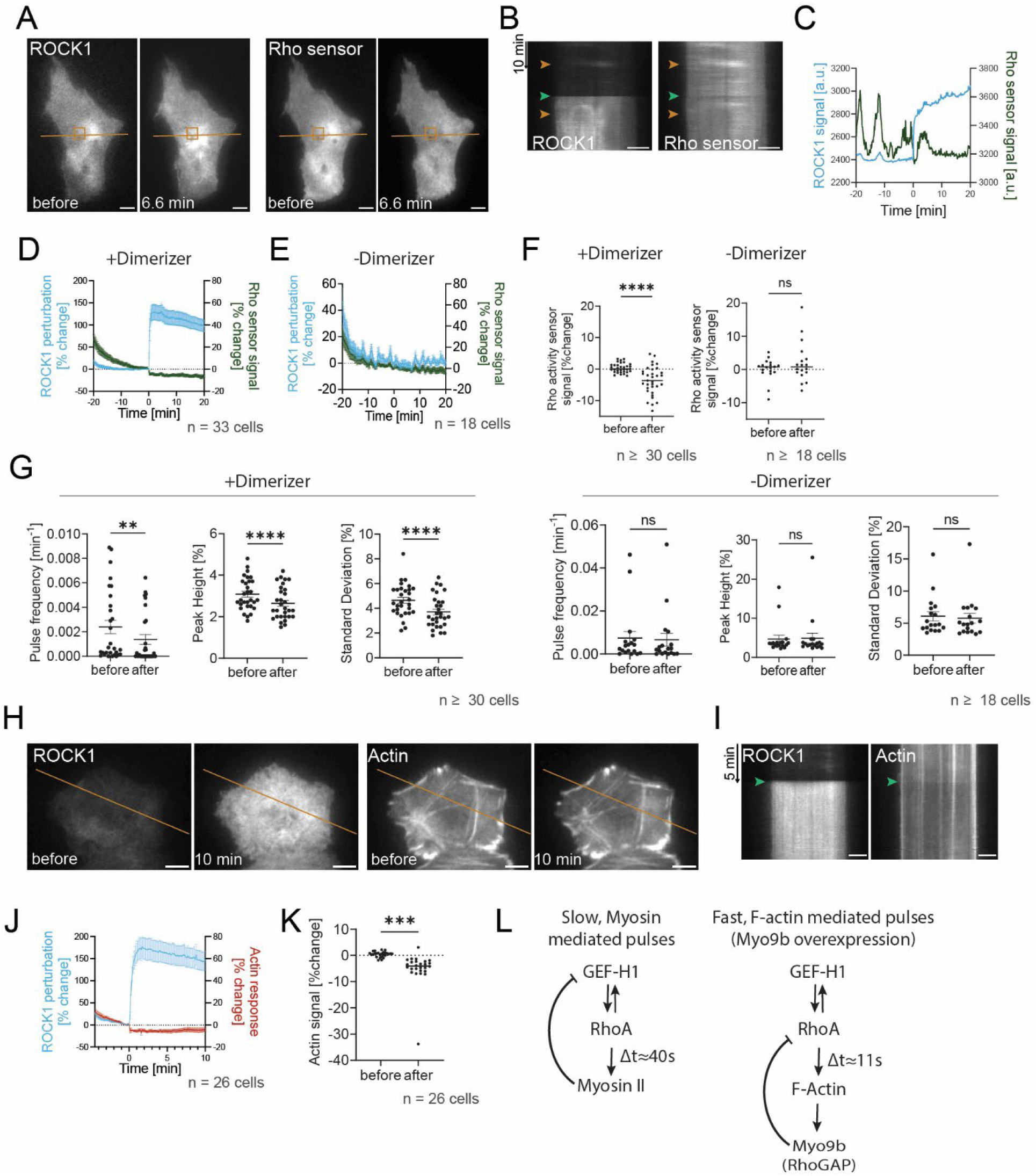
Rapid inhibition of cell contraction signal network dynamics after light-controlled activation of Myosin IIa. A-G: Analysis of Rhotekin-based mCherry-Rho activity sensor response to ROCK1 recruitment to the plasma membrane. A: Representative TIRF images of chemo-optogenetic mTurquoise2-ROCK1-eDHFR and mCherry-Rho sensor response. B: Kymographs corresponding to orange line in A. Orange arrowheads indicate time of the example frames in A. The green arrowhead indicates the timepoint of ROCK1 plasma membrane targeting, which precisely coincides with an immediate drop in basal Rho activity levels. C: Intensity profile of mTurqoise2-ROCK1-eDHFR and mCherry Rho activity sensor of the orange box in A. D-E: Quantification of mTurqoise-ROCK1-eDHFR perturbation and mCherry-Rho sensor response of cells treated with (D) or without (E) dimerizer (n ≥ 18 cells from 3 independent experiments). F: Quantification of the change in mCherry-Rho sensor fluorescence intensity at 3 time points before vs. 3 time points after perturbation. G: Quantification of mCherry-Rho sensor signal dynamics before and after induction of the light-based perturbation (+dimerizer: n ≥ 29 cells; –dimerizer: n ≥ 18 cells from 3 independent experiments). H-K: Analysis of Actin response to ROCK1 recruitment to the plasma membrane. H: Representative TIRF images of chemo-optogenetic mTurquoise2-ROCK1-eDHFR plasma membrane recruitment and the mCherry-Actin response. I: Kymographs corresponding to orange lines in H. The green arrowheads indicate the timepoint, at which ROCK1 plasma membrane targeting was triggered. J: Quantification of mCherry-actin reduction and mTurqoise-ROCK1-eDHFR perturbation (n = 26 cells from 3 independent experiments). K: Quantification of the change in mCherry-actin fluorescence intensity at 3 time points before vs. 3 time points after perturbation. L: Topology of the cell contraction signaling network in adherend mammalian cells. Left: Topology of the cell contraction signal network in adherent cells, in which Myosin II or Myosin II associated components mediate negative feedback regulation, for example via the inhibition of GEF-H1 by Myosin II (Lee et al., 2010). Right: Faster, F-Actin associated negative feedback that can be enforced in adherent mammalian cells (Graessl et al., 2017). Similar mechanisms might operate in *Xenopus* or starfish oozytes or *C. elegans* embryos. A-K: In all experiments, cells were treated with nocodazole (30 µM) to stimulate cell contraction dynamics. Error bars represent standard error of the mean. **: p < 0.01, ***: p < 0.001, ****: p < 0.0001, paired Student’s t-Test. Scale bars: 10 µm.

To investigate, if actin might play a role in the observed inhibitory action of ROCK1 at the plasma membrane, we directly measured actin levels at the plasma membrane (Figure 5H-I). Notably, actin levels did not increase but instead were even reduced by about 5% immediately upon our ROCK1 perturbation and stayed at this lower level for the subsequent 10 minutes (Figure 5J-K). These results provide definitive evidence that Myosin or a Myosin associated molecule, and not actin, mediates negative feedback regulation in the cell contraction signal network of adherent, mammalian cells.

Our observation that Myosin II acts as the negative feedback mediator is particularly striking, as it shows that adherent mammalian cells utilize a distinct molecular mechanism to generate cell contraction pulses compared to *Xenopus* or Starfish oozytes, or *C. elegans* embryos. Although all these systems generate activity patterns that are very similar, including pulses, oscillations and propagating waves, it should be noted that several specific features are also quite distinct. For example, adherent mammalian cells utilize Lbc-type RhoGEFs, such as GEF-H1 or LARG as positive feedback mediators, while oozytes rely on the non-Lbc type GEF Ect2 (Bement et al., 2015), which is specifically activated during mitosis (Tatsumoto et al., 1999). Furthermore, the period of signal network activity pulses is considerably longer for the Myosin-dependent mechanism in adherent mammalian cells (≈4min) (Kamps et al., 2020), compared to the Myosin-independent mechanisms in other systems (*C.elegans* embryos: period ≈0.5min (Nishikawa et al., 2017); *Xenopus* oozytes: period ≈1.5min (Michaud et al., 2022); Starfish oozytes: period ≈1.2min (Bement et al., 2015)). Notably, enforcing an alternative, actin-dependent negative feedback mechanism in adherent mammalian cells by overexpression of the F-actin associated RhoGAP Myo9b also leads to much shorter signal network activity pulses (period ≈0.5min (Graessl et al., 2017), Figure 5L), which more closely resemble the pulse periods in *C. elegans* embryos or *Xenopus* and starfish oozytes. These differences in pulse frequency for Myosin-dependent or actin-dependent negative feedback mechanisms are likely due to the differences in recruitment kinetics of Myosin II vs. actin downstream of Rho activity, which differ substantially in adherent mammalian cells (Myosin II: ≈40s; Actin ≈11s (Graessl et al., 2017)).

We were also surprised by the observation that CA-ROCK1 can shift pulsatile Rho activity dynamics away from central cell attachment areas and restrict these pulses at the cell periphery. Our previous studies suggest that central Rho activity pulses play a role in mechanotransduction in non-migratory U2OS cells (Graessl et al., 2017), while peripheral Rho activity pulses are typically associated with protrusion retraction cycles, that are often observed in highly migratory cells, such as the A431 epidermal carcinoma cell line (Nanda et al., 2023). While CA-ROCK1 overexpression constitutively activates Myosin in a highly artificial fashion, Rho independent mechanisms that can activate ROCK1 have been described previously, for example in apoptosis (Coleman et al., 2001; Sebbagh et al., 2001) and in thrombin-induced endothelial microparticle generation (Sapet et al., 2006), and might have a similar effect on the spatio-temporal organization of Rho activity patterns.

In conclusion, we show that adherent mammalian cells utilize a specific signal network to control cell contraction dynamics, which is dependent on a slow, Myosin-II mediated negative feedback loop (Figure 5L).

## Methods

### Cell Culture

U2OS cells were cultivated in Dulbecco’s modified Eagle medium (DMEM) with GlutaMAX™ supplement (Gibco, Cat. No. 31966-047), 10% fetal bovine serum (FBS) (Gibco, Cat. No. A5256701) and 10 units/ml penicillin and 100 µg/ml streptomycin (PAN Biotech, Cat. No. P06-07100). The maintenance of cells was carried out in accordance to standard culture techniques, employing Trypsin/EDTA (0.05%/0.02%, PAN Biotech, Cat. No.: P10-02355P, TC dishes Cell+ Sarstedt) at 37°C and 5% CO2. For standard live cell imaging LabTek glass surface slides (Thermo Fisher Scientific) were coated with 10 µg/ml calf skin collagen-I (Sigma C8919) and cells were plated onto these slides one day prior to transfection. The transfection of all plasmids into U2OS cells was conducted using Lipofectamine™ 2000 (Thermo Fisher Scientific,

Cat. No. 11668019). Live-cell imaging was performed in CO₂-independent, HEPES-stabilized imaging medium (Pan Biotech, Cat. No. P04-01163), supplemented with 10% FBS, at 34 °C, at the indicated frame rates.

### Plasmid construction

CMV-mCitrine-GEF-H1 C53R was generated via a 2 fragment Gibson assembly, using EGFP-GEF-H1 C53R (Krendel et al., 2002) as the backbone after removing the EGFP coding sequence using the EcoRI and KpnI and a mCitrine insert obtained from delCMV-mCitrine (Nanda et al., 2023) using 5’-ctcgtttagtgaaccgtcagaattcgccaccatggtgagcaagggcgag-3’ as forward and 5’-ttcgatccgagacatggtacccttgtacagctcgtccatgc-3’ as reverse primers. The full length ROCK1 expression construct mCherry-ROCK1^1^ was generated via Gateway cloning using the entry vector R777-E283 Hs.ROCK1 (Addgene plasmid #70567) and destination vector pDEST-CMV-N-mCherry (Addgene plasmid #123215) (Agrotis et al., 2019). The constitutively active mutant mCherry-CA-ROCK1 was derived from mCherry-ROCK1 by truncation of the C-terminal sequence 478-1354 via a 1 fragment Gibson assembly using 5’-gaaatcaaagaagaaatctaTAATCAACTTTCTTGTACAAAGTG-3’ as forward primer and 5’-TAGATTTCTTCTTTGATTTCCC-3’ as reverse primer. The plasmid mTurqoise-MyosinIIA was obtained from Addgene (plasmid #55571) and the generation of mCherry-NMHCIIA was already described in Kowalczyk et al. 2022. mTurquoise-ROCK1-eDHFR was generated via a 2 fragment Gibson assembly, using mTurquoise2-eDHFR (Kamps et al., 2020) cut with XhoI as the backbone and a ROCK1 insert that was amplified from mCherry-ROCK1 using 5’-ctggactcgtacaagatatcaatgtcgactggggacagttttg-3’ and 5’-gcaatcagactgatcatagccgatccacttccagaaccggaactagtttttccagatgtatttttgacc-3’ as primers. The artificial receptor for MAP experiments, CMV-Dimerizer-PARC-CCL-mCitrine-Halotag, was described previously (Chen et al. 2017). The Rhotekin-based Rho sensors, delCMV-mCherry-RBD and delCMV-mCitrine-RBD, were both already described in Graessl et al. (2017) and Kamps et al. (2020), respectively. Lastly, mCherry-actin was already described in Nanda et al. (2023). As control for mCherry-CA-ROCK1, the empty Gateway cloning destination vector pDEST-CMV-N-mCherry from Addgene (#123215) was used.

### Microscopy

Total internal reflection fluorescence (TIRF) microscopy was performed on an Olympus IX-81 microscope equipped with a TIRF-MITICO motorized TIRF illumination combiner, an Apo TIRF 60× 1.45 NA oil immersion objective, and a ZDC autofocus device. Four lasers were used for TIRF illumination: Cell R diode lasers (Olympus) with wavelengths of 405 nm (50 mW), 445 nm (50 mW), and 561 nm (100 mW) and a 514-nm OBIS diode laser (150 mW; Coherent, Inc., Santa Clara, CA, USA). Additionally, the Spectra X light engine (Lumencor) was used for wide-field illumination. The C9100-13 EMCCD camera (Hamamatsu, Herrsching am Ammersee, Germany) without binning at medium gain was used for signal detection.

### Fixed Cell Imaging

Cells were fixed with 4% paraformaldehyde (PFA) for 15 minutes at 18-22 h after transfection. Afterwards, cells were stained with 1:1000 phalloidin-Atto 488 (Sigma-Aldrich, 49409) and 1:1000 Hoechst 33342 (Sigma-Adrich) overnight and imaged using wide-field illumination.

### Molecular Activity Painting (MAP)

Molecular activity painting was described in detail previously in Chen et al. (2017), and the simplified method used in this work was previously described in Kamps et al. (2020). The illumination in a diffraction-limited spot was performed for 400 ms at 180 nW (laser power measured at the aperture of the 60x objective). Here, in addition, a new variant was developed to introduce perturbations in the entire cell attachment area. First, 17 000 U2OS cells were plated onto Labtek dishes and transfected as described above. Incubation with Nvoc-TMP-Cl photocaged chemical dimerizer and washing steps were identical to the protocol described in Kamps et al. (2020). Photo uncaging in the entire cell attachment area was achieved using a 405 nm laser for 400 ms in TIRF mode of the TIR-MITICO motorized TIRF illumination combiner (Olympus). To compensate for the lower illumination density in TIRF illumination compared to a diffraction-limied spot, 3x higher laser power was used for the new molecular activity painting variant to uncage the dimerizer in the entire cell attachment area.

### Image and Video Analysis

All image stacks were prepared for later automated or manual analysis using custom made image J macros. The ImageJ plugin *Image Stabilizer* was used to stabilize the field of view in one channel and the stabilization log generated by this plugin was applied to the other channel(s) using the *Image Stabilizer log applier* (Kang Li, 2009).

Evaluation of myosin structures (i.e. the presence of a single myosin focus vs normal stress fibers or multiple small patches of Myosin) and the qualitative evaluation of GEF-H1 spatio-temporal activity patterns (peripheral activity pulses vs central or no activity pulse) were carried out in a single-blind design. This was achieved by the imageJ plug in *Analyse & Decide* which enabled the presentation of videos in an arbitrary sequence to an individual who was blinded to the treatment assignments.

All statistical analyses were performed in *Prism10*.

#### Quantitative analysis of pulsatory signal network dynamics

The analysis of signal network dynamics (e.g. peak height, peak frequency and local signal standard deviation) was performed essentially as described previously (Graessl et al., 2017). Briefly, cells were imaged every 8 seconds for a total duration of 30 min. In the recorded image sequences, individual cells were isolated manually and scaled down by a factor of 10×10 pixels to reduce noise. Thresholds were identified manually to define the borders of the scaled-down cells. The automated Peak analysis script previously described in Graessl et al. (2017) was modified to facilitate the transfer of cell masks between different channels.

#### Analysis of the response to chemo-optogenetic perturbations

To measure the response to local chemo-optogenetic perturbations, the perturbation spot was selected. Next, the fluorescence intensity in the perturbation and response channel were measured as a function of time and normalized to the time point prior to the perturbation. The normalized data was then plotted in *Prism10* to represent the perturbation– and response strength. Measurements were only included in subsequent analysis, if substantial perturbations with a signal increase of at least 5% were achieved.

The response to chemo-optogenetic perturbations in the whole cell attachment area was analyzed using a modification of the “Quantitative analysis of pulsatory signal network dynamics” described above. Briefly, individual cells were isolated manually, and the fluorescence intensity was measured in the whole cell. This was performed using a custom-made *imageJ* macro that automatically detects the cell boundaries in a selected channel and measures the fluorescence intensity successively in all channels. Here, the readout channel (i.e. Myosin, Rho or Actin) was used as it was best suited to define the cell borders. Subsequently, the measured intensities were normalized to the time point prior to perturbation. To compare the cell contraction network dynamics before and after perturbation, the image sequences were separated into prior and after perturbation and the pulse analysis was carried out as described above for each individualized cell in each stack.

### Numerical simulations

All simulations that were performed in this study were based on a system of stochastic differential equations (SDE) and parameters that were derived in a previous study(Kamps et al., 2020). Briefly, the system comprises three variables G, R, and M which correspond to the active form of GEF-H1, Rho, and Myosin, respectively, as a function of time t:

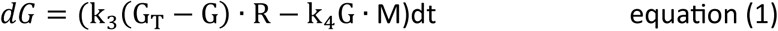

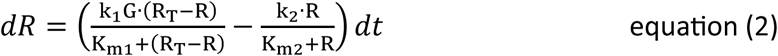

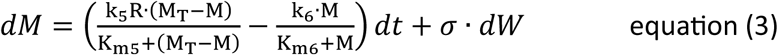

G_T_, R_T_, M_T_ are the total concentrations of GEF-H1, Rho, and Myosin, that are assumed to be constant over time. k_1_-k_6_ are rate constants and K_m1_, K_m2,_ K_m5,_ K_m6_ represent Michaelis constants and σ represents the level of gaussian noise. Here, we extended this previously published SDE system and included an additional Michaelis-Menten term with rate constant k_7_ and Michaelis constant K_m7_ that represents the Rho independent activation of Myosin by constitutively active ROCK:

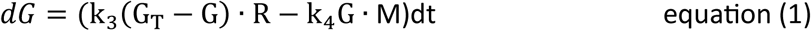

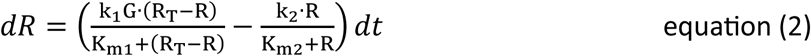

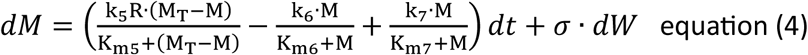

*with*: 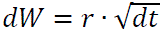

The reactions that form the basis of this system are shown schematically in Figure 2A. In our previous study(Kamps et al., 2020), we used data from the literature, our own experimental data using the same experimental system (i.e. U2OS cells expressing GEF-H1 C53R) and a Bayesian fitting approach to estimate the values for the parameters G_T_, R_T_, M_T_, k_1_-k_6_ and K_m1_, K_m2,_ K_m5,_ K_m6_, and we used these parameters here, in our current study, to perform simulations of the stimulated, unperturbed state of this system (i.e. with increased total GEF-H1 concentrations corresponding to G_T_=0.3 and without constitutively active ROCK corresponding to k_7_=0, and a level of gaussian noise of σ=0.001, shown in Figure 2B, left).

Simulations that predict the effect of constitutively active ROCK1 expression (Figure 2B-D) included increasing values for k_7_. Similarly, simulations that predict the effect of Myosin inhibition via blebbistatin (Figure 2B and Figure S1A-B) or of ROCK inhibition via Y27632 (Figure 2B and Figure S1C-D) included decreasing values for M_T_ or k_5_, respectively. The exact values of k_7_, or the percent reduction of the values for M_T_ or k_5_, are indicated in the corresponding Figure panels or the corresponding x-axes of plots. For K_m7_ we did not have previous estimations from our studies or literature and did not assume a specific value and therefore we set this parameter to equal 1.

## Supporting information

Movie_Figure3A_left

Movie_Figure3A_right

Movie_Figure3D_left

Movie_Figure3D_middle

Movie_FigureD_right

## Acknowledgements

We would like to thank Dominic Kamps for generating the mTurquoise2-NES-Linker-eDHFR construct, which was used for further cloning. We are thankful for helpful discussions with Perihan Nalbant, Peter Bieling and Rudolf Merkel and Ricarda Lüttig for assistance in data analysis. This work was supported by the Deutsche Forschungsgesellschaft DFG Heisenberg Programme grant DE 823/8-1, the DFG project grant DE 823/9-1 and the DFG principal investigator grant DE 823/10-1 to L.D.

## Author contributions

L.D. and C.G. designed the research. C.G. performed and analyzed the majority of experiments and numerical simulations. J.H. performed experiments to study the long-term effects of CA-ROCK1 shown in Figure 2. L.D. supervised all experiments. C.G. and L.D. wrote the manuscript.

## Supplemental Material

**Figure S1:**
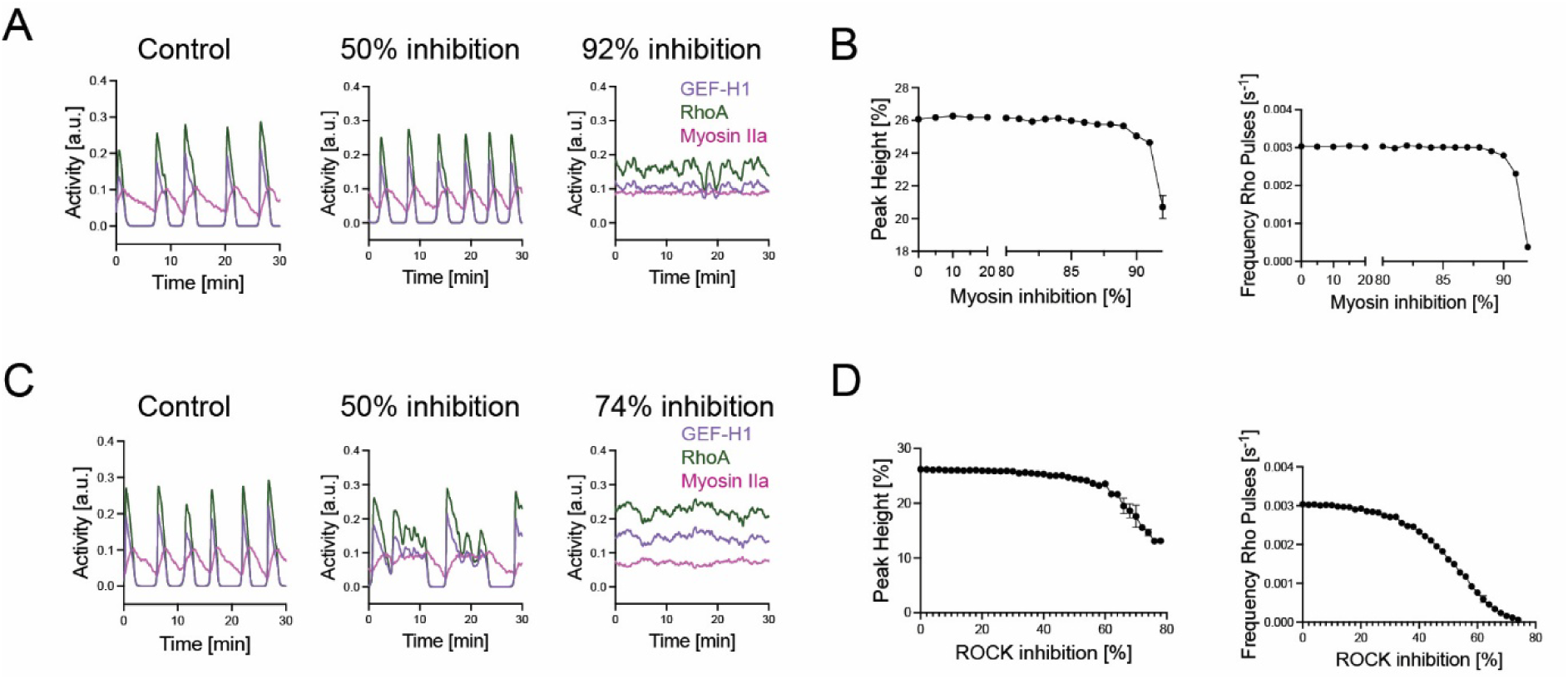
Theoretical investigation of the effect of Myosin or ROCK inhibition on dynamics of the cell contraction signal network. A: SDE simulations of the cell contraction signal network activity dynamics with indicated levels of Myosin inhibition. B: Dependence of Rho activity peak height (left) and Rho pulse frequency (right) on increasing levels of Myosin inhibition. Myosin inhibition in the x-axis corresponds to the % reduction of the parameter MT that represents the total myosin concentration (i.e. the sum of active and inactive Myosin in the scheme shown in Figure 2A). C: SDE simulations of the cell contraction signal network activity dynamics with indicated levels of ROCK inhibition. D: Dependence of Rho activity peak heigt (left) and Rho pulse frequency (right) on increasing levels of ROCK inhibition. ROCK inhibition in the x-axis corresponds to the % reduction of the parameter k5 in the scheme shown in Figure 2A, which represents the rate of the reaction that activates Myosin downstream of active Rho. Error bars represent standard error of the mean of 3 independent simulations each.

### Movie Legends

Movie_Figure1: Myosin IIa strongly correlates with GEF-H1 C53R stimulated cell contraction signal network activity in space and time. Time-lapse TIRF videos of mCitrine-GEF-H1 C53R and mTurquoise-Myosin-IIa in a representative U2OS cell. Imaging started 42 h after plating on collagen-coated glass-bottom dishes. Images were collected with a frame rate of 7.5/min. Scale bars 10 µm.

Movie_Figure3A_left: Myosin forms highly dynamic cortical structure that are organized into stress fibers and local pulsatory patterns in the absence of CA-ROCK1. Time-lapse TIRF videos of a mCherry-based volume marker as control and mCitrine-Myosin-IIa in a representative U2OS cell. Imaging started 42 h after plating on collagen-coated glass-bottom dishes. Images were collected with a frame rate of 7.5/min. Scale bars 10 µm.

Movie_Figure3A_right: Myosin forms a single, dominant Myosin II focus via a contractile flow from the periphery towards the cell center in the presence of CA-ROCK1. Time-lapse TIRF videos of mCherry-CA-ROCK1 and mCitrine-Myosin-IIa in a representative U2OS cell. Imaging started 42 h after plating on collagen-coated glass-bottom dishes. Images were collected with a frame rate of 7.5/min. Scale bars 10 µm.

Movie_Figure3D_left: Pulse– and wave-like plasma membrane recrutiment patterns of GEF-H1 C53R in the absence of CA-ROCK1. Time-lapse TIRF videos of mCherry-CA-ROCK1 and mCitrine-GEF-H1 C53R in a representative U2OS cell. Imaging started 42 h after plating on collagen-coated glass-bottom dishes. Images were collected with a frame rate of 7.5/min. Scale bars 10 µm.

Movie_Figure3D_middle: In a subset of cells, plasma membrane recrutiment patterns of GEF-H1 C53R dynamics are shifted towards the periphery of the cell attachment area in presence of CA-ROCK1. Time-lapse TIRF videos of mCherry-CA-ROCK1 and mCitrine-GEF-H1 C53R in a representative U2OS cell. Imaging started 42 h after plating on collagen-coated glass-bottom dishes. Images were collected with a frame rate of 7.5/min. Scale bars 10 µm.

Movie_Figure3D_right: In a subset of cells, pulse– and wave-like plasma membrane recruitment patterns of GEF-H1 C53R are abolished in presence of CA-ROCK1. Time-lapse TIRF videos of mCherry-CA-ROCK1 and mCitrine-GEF-H1 C53R in a representative U2OS cell. Imaging started 42 h after plating on collagen-coated glass-bottom dishes. Images were collected with a frame rate of 7.5/min. Scale bars 10 µm.

Movie_Figure3H-I: Spatio-temporal separation between local GEF-H1 C53R activity pulses and a Myosin II focus in the presence of CA-ROCK1. Time-lapse TIRF video of mCitrine-GEF-H1 C53R and mTurquoise-Myosin-IIa in a representative U2OS cell. Imaging started 42 h after plating on collagen-coated glass-bottom dishes. Images were collected with a frame rate of 7.5/min. Scale bars 10 µm.

Movie_Figure4B-C: Rapid chemo-optogenetic plasma membrane recruitment of ROCK1 activates and recruits Myosin II at a distinct site at the plasma membrane. TIRF video-microscopy of a representative U2OS cell, expressing the perturbation construct mTurquoise2-ROCK1-eDHFR (left), the response construct mCherry-non-muscle Myosin heavy chain IIa and the artificial receptor mCitrine-VSVG HaloTag-PARC. A focused light pulse at 405nm was applied at timepoint 0 to target the perturbation construct to a small region of the plasma membrane. Images were collected with a frame rate of 6/min. Scale bars 10 µm.

Movie_Figure4G-H: Rapid chemo-optogenetic plasma membrane recruitment of ROCK1 activates and recruits Myosin II. TIRF video-microscopy of a representative U2OS cell, expressing the perturbation construct mTurquoise2-ROCK1-eDHFR (left), the response construct mCherry-non-muscle Myosin heavy chain IIa and the artificial receptor mCitrine-VSVG HaloTag-PARC. Photo-uncaging in the entire cell attachement area via TIRF illumination was applied at timepoint 0 to target the perturbation construct to the plasma membrane receptor. Images were collected with a frame rate of 7.5/min. Scale bars 10 µm.

Movie_Figure5A-B: Rapid chemo-optogenetic plasma membrane recruitment of ROCK1 reduces Rho activity at the plasma membrane. TIRF video-microscopy of a representative U2OS cell, expressing the perturbation construct mTurquoise2-ROCK1-eDHFR (left), the Rhotekin-based Rho activity sensor as response construct (right) and the artificial receptor mCitrine-VSVG HaloTag-PARC. Photo-uncaging in the entire cell attachement area via TIRF illumination was applied at timepoint 0 to target the perturbation construct to the plasma membrane receptor. Images were collected with a frame rate of 7.5/min. Scale bars 10 µm.

Movie_Figure5H-I: Rapid chemo-optogenetic plasma membrane recruitment of ROCK1 reduces actin localization at the plasma membrane. TIRF video-microscopy of a representative U2OS cell expressing the perturbation construct mTurquoise2-ROCK1-eDHFR (left), the response construct mCherry-Actin-UB (mCherry-Actin fusion protein with a long linker between the fusion partners and driven by the Ubiquitin C promotor) and the artificial receptor mCitrine-VSVG HaloTag-PARC. Photo-uncaging in the entire cell attachement area via TIRF illumination was applied at timepoint 0 to target the perturbation construct to the plasma membrane receptor. Images were collected with a frame rate of 7.5/min. Scale bars 10 µm.

## Footnotes

1 This construct was primarily developed and generated in a companion study by Gierse et al. 2025 that is submitted together with this work. The experimental details for the construction of this construct are briefly stated in both papers. For future derivative work that use mCherry-ROCK1, the companion study should be cited.

